# Visual temporal integration by multi-level regularities fosters the emergence of dynamic conscious experience

**DOI:** 10.1101/2021.04.28.440365

**Authors:** Ruichen Hu, Peijun Yuan, Ying Wang, Yi Jiang

**Affiliations:** State Key Laboratory of Brain and Cognitive Science, CAS Center for Excellence in Brain Science and Intelligence Technology, Institute of Psychology, Chinese Academy of Sciences, 16 Lincui Road, Beijing 100101, P. R. China; Department of Psychology, University of Chinese Academy of Sciences, 19A Yuquan Road, Beijing 100049, P. R. China

## Abstract

The relationship between information integration and visual awareness is central to contemporary theories and research on human consciousness. While there is evidence that humans are adept at integrating spatially structured information to form a coherent conscious percept, to date, little is known about how we integrate visual information over time based on its temporal structure and whether such temporal integration process contributes to our awareness of the dynamic world. Using binocular rivalry, we demonstrated that a diverse set of structured visual streams, constituted either by idiom, shape, or motion stimuli, predominated over their non-structured but otherwise matched counterparts in the competition for conscious access. Despite the apparent resemblance, there was a substantial dissociation of these observed privileges between the semantic- and perceptual-level structures, specifically regarding their resistance to spatiotemporal perturbations and demands for conscious processing over the visual integration process. These findings corroborate the essential role of structure-guided information integration in the generation of conscious content and highlight temporal integration by multi-level regularities as a fundamental mechanism to foster the emergence of continuous conscious experience.

## Introduction

With a simple glance at Van Gogh’s Starry Night, you would be impressed by a deep blue sky roiling with shining stars above a tranquil village, all as a meaningful whole rather than a collection of unrelated attributes. As you walk through the gallery, this particular conscious experience could be extended both across space and over time, engendering a coherent and continuous awareness of the external world. In this regard, the emergence of visual consciousness entails the ability to bind different features into impartible objects (Engel et al., 1999; Singer, 2001) and integrate disparate elements into a unitary conceptual structure across multiple spatial and temporal scales (Hoel et al., 2016; Tononi and Koch, 2015). Accordingly, intelligent creatures who can generate highly composite conceptual structures, presumably via incorporating intrinsic structures of the external inputs based on the underlying regularities, may gain an adaptive advantage to live in a complex environment (Tononi et al., 2016).

Humans are endowed with a remarkable capability to extract regularities embedded in the spatial structure of their visual environment for the construction of conscious perception. The evidence dates back to the early twentieth century when psychologists found people follow a set of Gestalt laws to group spatially non-overlapped visual elements into perceptually meaningful entities (Koffka, 1935). In contrast, only in recent decades, there has been an emerging interest in how human beings utilize regularities from the temporal structure of the incoming visual information to promote conscious perception and guide adaptive behavior. Strikingly, even in the absence of configurational cues, temporal structures defined merely by synchronous changes in a visual feature can lead to immediate perception of dynamic objects from a mixture of elements (Alais et al., 1998; Lee and Blake, 1999). Also, temporal structures based on statistical regularities can attract attention to specific visual features or spatial locations (Zhao et al., 2013). Furthermore, complex temporal structures in sign languages, similar to that in auditory speech and music (Ding et al., 2016; Doelling and Poeppel, 2015), can entrain neural oscillations in the visual cortex to maximize sensitivity to rhythmic visual information (Brookshire et al., 2017).

The existing research shows that temporal structures defined by various types of regularities may influence our conscious experience of the dynamic world. Nevertheless, whether regular temporal structure confers a direct benefit to the emergence of conscious content in the continuum of time remains undetermined. The answer to this question is of considerable importance because the content of consciousness is extremely limited, in contrast to the seemingly unlimited information available in the visual environment (Cohen and Dennett, 2011; Kouider et al., 2010). To resolve this paradox, the visual system has to prioritize certain types of content (e.g., those physically or biologically salient stimuli) over the others during their competition for conscious access (Alpers and Gerdes, 2007; Yu and Blake, 1992). Despite its significance, our knowledge about the rules and mechanisms governing such visual competition is far from complete, especially in the time domain (Brascamp et al., 2018). Here we proposed that regularities in the temporal dimension could be a fundamental factor underlying the construction of visual awareness in a dynamic environment. In particular, a highly regular temporal structure may facilitate spatiotemporal integration of the incoming visual information, which is taken as a prerequisite for the construction of coherent conscious experiences (Hoel et al., 2016; Tononi et al., 2016), thereby empowering a visual stream to stand out from others during the visual competition.

In a broad sense, a structured information stream may consist of stimuli organized in time based on their perceptual properties (Henry et al., 2014) or symbolic meanings (Ding et al., 2016; Doelling and Poeppel, 2015). Correspondingly, the temporal structures in dynamic visual information may derive from regularities at different levels (e.g., perceptual and semantic), which brings up another important question: whether the influence of temporal structure on visual competition, if observed, can extend across information levels. To address this issue, we assessed the potential advantages of structured streams derived either from perceptual-level or semantic-level regularity, relative to their non-structured counterparts. More importantly, to help elucidate the mechanisms responsible for the observed advantages, we further characterized the effects driven by these multi-level regularities from the perspective of temporal information integration.

## Results

### Visual awareness is biased towards temporally structured information

In Experiment 1, we set out to assess whether temporal structures defined by different levels of regularities could modulate conscious awareness in binocular rivalry (BR), a phenomenon widely used to examine how a conscious mind resolves the conflict between two competing visual representations throughout different stages of visual processing (Blake and Logothetis, 2002). During BR, with two distinct stimuli presented to their left and right eyes, observers experience spontaneous alternations between two conscious percepts. The relative dominance duration of these perceptual outcomes can serve as an index to determine whether a visual stimulus with a particular property outperforms its competitor in terms of perceptual predominance. We pitted structured streams composed of periodically changing stimuli regarding their shape, motion, or contrast (perceptual-level regularity) and those of concatenated Chinese idioms (semantic-level regularity) against their physically matched but temporally randomized counterparts (Fig. 1a). If dynamic information with regular temporal structure enjoys a privilege in the visual competition, we would observe prolonged dominance durations (or a dominance ratio larger than 1, see Methods for details) for the structured stream relative to the random one.

**Fig. 1.**
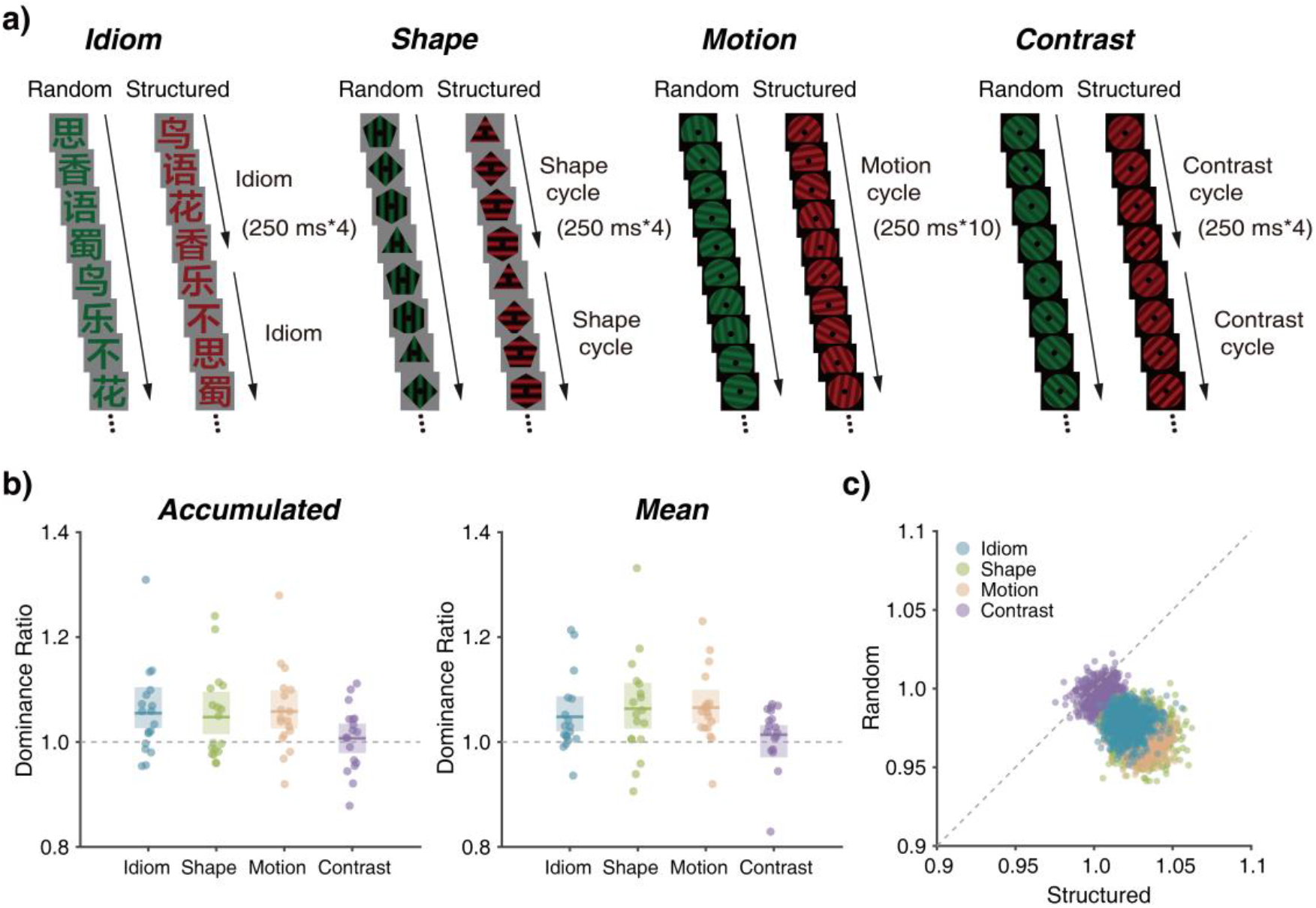
Example stimuli and results of Experiment 1. a) For each stimulus type, structured and random streams composed of identical elements but with different stimulus sequences were presented dichoptically to participants. b) The temporal structure advantage for idiom, shape, and motion, but not for contrast, as indicated by the dominance ratio indices. Dominance ratios were obtained by dividing the accumulated (trial-based) and mean (percept-based) dominance durations of the structured streams with that of the random streams. A dominance ratio larger than 1 reveals a relative predominance of the structure stream. The colored lines and bars represent the group means with bootstrapped 95% CIs, and each colored dot represents data from an individual participant. c) Bootstrap distributions of mean normalized dominance durations show a dissociation between the contrast and the other stimulus conditions.

In accordance with our assumption, one sample t-tests revealed an accumulated dominance ratio significantly greater than 1 for the idiom streams (Fig. 1b; *t* (17) = 2.78, *p* = 0.01, Cohen’ *d* = 0.65, two-tailed, the same below), suggesting that semantic-level regularities can promote awareness dominance during the visual competition. Similar patterns were observed for perceptual-level regularities in the shape and motion conditions (Shape: *t* (17) = 2.40, *p* = 0.02, Cohen’ *d* = 0.56; Motion: *t* (17) = 3.06, *p* < 0.01, Cohen’ *d* = 0.72), while not in the contrast condition (*t* (17) = 0.48, *p* = 0.63, Cohen’ *d* = 0.11). Consistent with these findings, analyses on the mean dominance ratios yielded similar results: the temporal structure advantage was found in the idiom (*t* (17) = 2.80, *p* = 0.01, Cohen’ *d* = 0.66), shape (*t* (17) = 2.77, *p* = 0.01, Cohen’ *d* = 0.65), and motion (*t* (17) = 3.96, *p* < 0.001, Cohen’ *d* = 0.93) conditions, but not in the contrast condition (*t* (17) = 1.01, *p* = 0.32, Cohen’ *d* = 0.23).

To verify the differences among stimulus conditions, one thousand bootstrapped samples of the normalized dominance durations for each stimulus condition were shown in Fig. 1c, with the horizontal axis representing the structured streams and the vertical axis representing the random streams. While the distribution of the contrast stimulus is clustered around and nearly bisected by the diagonal line (y=x), clusters of the other stimuli all fall below this line, confirming that only in the later conditions, the dominance durations are reliably prolonged by structured visual information.

### Dissociation between semantic- and perceptual-level regularities: the resistance to spatiotemporal perturbations

We have shown that temporal structure based on regularities at the semantic (as in the idiom condition) and the perceptual (as in the shape and motion conditions) levels both enjoyed prolonged dominance durations during visual competition. These findings leave open the question of whether such cross-level advantage is driven by a common mechanism. As aforementioned, spatiotemporal integration of isolated visual cues is key to the emergence of visual awareness (Hoel et al., 2016; Tononi et al., 2016). Here we asked whether the privilege enjoyed by different levels of regularities would be constrained, to a similar extent, by the uniformity of the spatiotemporal integration window.

We disrupted the uniform spatiotemporal arrangement of individual items in the rivalry streams by adding temporal (Experiment 2) or spatial (Experiment 3) jitters. In this way, we varied the temporal or spatial properties of the integration window without changing the information of the rivalry streams (Fig. 2a and 2b).

**Fig. 2.**
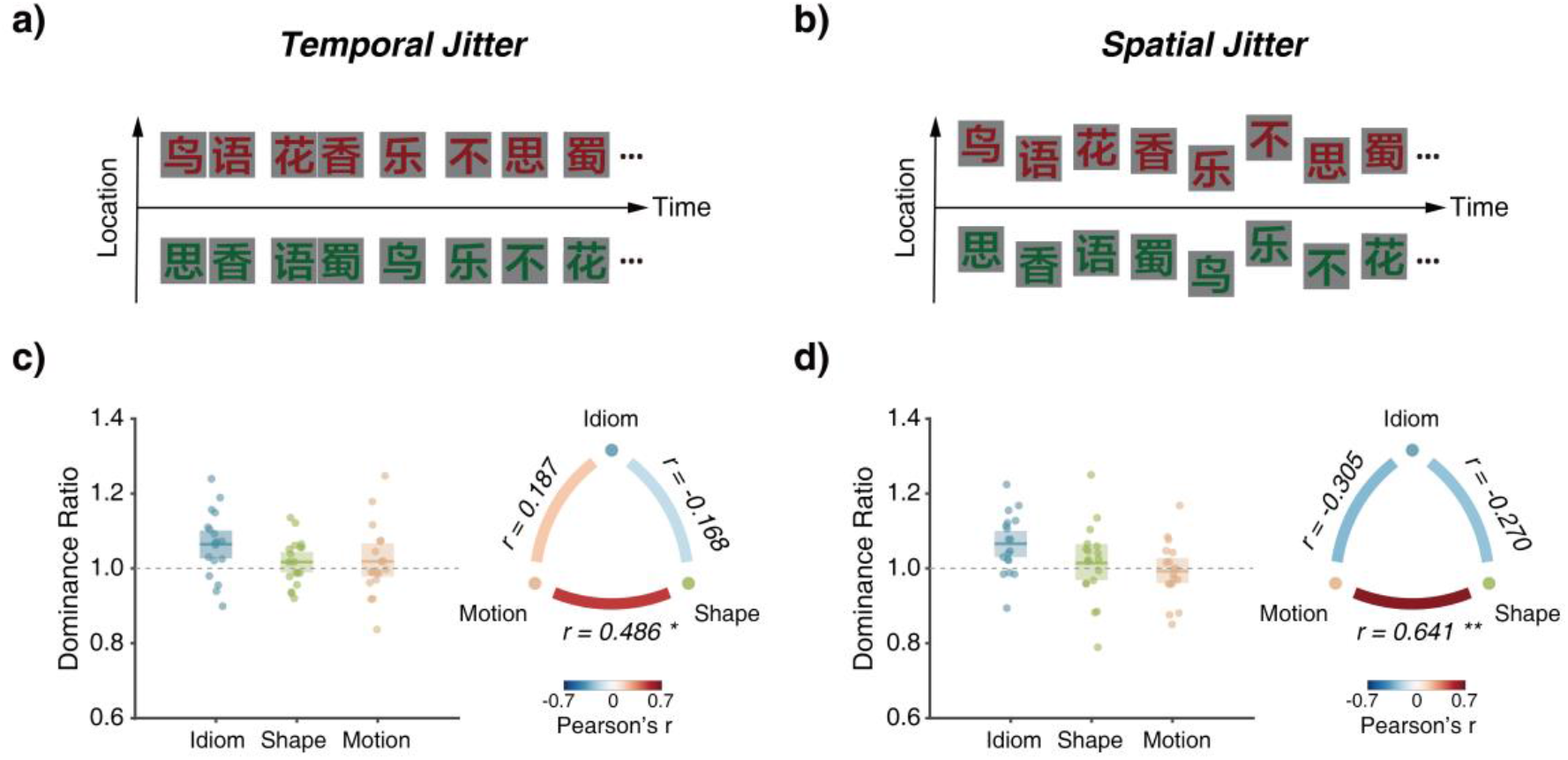
Illustrations of experimental manipulations and results of Experiments 2 and 3. Stimulus streams used in Experiments 2 and 3 were the same as that in the idiom, shape, and motion conditions of Experiment 1, except that a) the temporal duration of each stimulus in Experiment 2 was not constant, but ranged randomly from 125 ms to 375 ms; and b) the spatial location of each stimulus in Experiment 3 was not fixed, but varied randomly from 0° to 1.51° to the retinal center. Note that the spatiotemporal perturbations were illustrated on the idiom stream only, while the same manipulations were also applied to the shape and motion streams. c) & d) Dominance ratios of the three stimulus conditions and the pairwise Pearson correlation results for Experiments 2 and 3. The colored lines and bars represent the averaged dominance ratios with bootstrapped 95% CIs, and each colored dot represents one participant. * p < 0.05; **: p < 0.01.

In the temporal jitter experiment, the dominance ratio of structured idiom streams over the random counterparts remained significantly greater than 1 (Fig. 2c; *t* (17) = 3.18, *p* < 0.01, Cohen’ *d* = 0.74), with the effect comparable to that observed with the isochronous sequences, whereas for the shape and motion conditions, the advantage of structured streams over the random ones no longer existed (Fig. 2c; Shape: *t* (17) = 1.18, *p* = 0.25, Cohen’ *d* = 0.27; Motion: *t* (17) = 0.81, *p* = 0.42, Cohen’ *d* = 0.19). Similar results were observed in the spatial jitter experiment. While the structured idiom streams had a dominance ratio significantly larger than 1(Fig. 2d; *t* (17) = 3.48, *p* < 0.01, Cohen’ *d* = 0.82), other stimulus conditions revealed no such tendency (Fig. 2d; Shape: *t* (17) = 0.58, *p* = 0.56, Cohen’ *d* = 0.13; Motion: *t* (17) = 0.41, *p* = 0.68, Cohen’ *d* = 0.09).

To quantify the association of the observed effects across stimulus types, we calculated the pairwise Pearson correlation coefficients of dominance ratios based on the mean dominance durations among the three stimulus conditions (Fig. 2c & 2d). For both Experiments 2 and 3, we observed a significant correlation, particularly within perceptual-level regularities, i.e., between the shape and motion conditions (Experiment 2: *r* = 0.49, *p* < 0.05; Experiment 3: *r* = 0.64, *p* < 0.01). By contrast, no significant correlations were observed across the perceptual and semantic levels, i.e., between the idiom and shape (Experiment 2: *r* = -0.17, *p* = 0.51; Experiment 3: *r* = -0.31, *p* = 0.22) or between the idiom and motion conditions (Experiment 2: *r* = 0.19, *p* = 0.46; Experiment 3: *r* = -0.27, *p* = 0.28).

Taken together, these results reveal a clear dissociation between the semantic- and perceptual-level regularities regarding the boundary conditions for them to modulate awareness. The finding that only the privilege of idiom streams was tolerant of spatiotemporal perturbations on the integration window points to the existence of a specialized mechanism for prioritizing structured semantic information during visual competitions, which is distinct from that underlying basic visual feature integration.

### Dissociation between semantic- and perceptual-level regularities: conscious and nonconscious benefits

The spatiotemporal integration of dynamic visual information may occur, at least to some extent, beyond awareness (Mudrik et al., 2014), which poses another fundamental question: whether the observed privileges of different types of regularities originate from the conscious or the nonconscious visual integration processes. Regularities in dynamic visual information may lengthen the dominance durations of the structured streams when they are consciously perceived, i.e., during the dominance phase of BR (Lee et al., 2015), or reduce the suppression durations of the structured streams when they are inhibited from awareness, i.e., during the dominance phase of the random streams (Blake, 1977; Lunghi and Alais, 2015), both of which can lead to a greater percentage of perceptual dominance. To distinguish between these possibilities, in Experiment 4, we added baseline conditions where the two rivalry streams were both deprived of regular temporal structures, but matched respectively with the structured and random streams in the experimental conditions to control for low-level factors (Fig. 3a & 3b). By comparing the experimental and baseline conditions, we expected to isolate potential effects pertinent to the dominance phase of the structured streams (conscious benefits) from those to their suppression phase (nonconscious benefits).

**Fig. 3.**
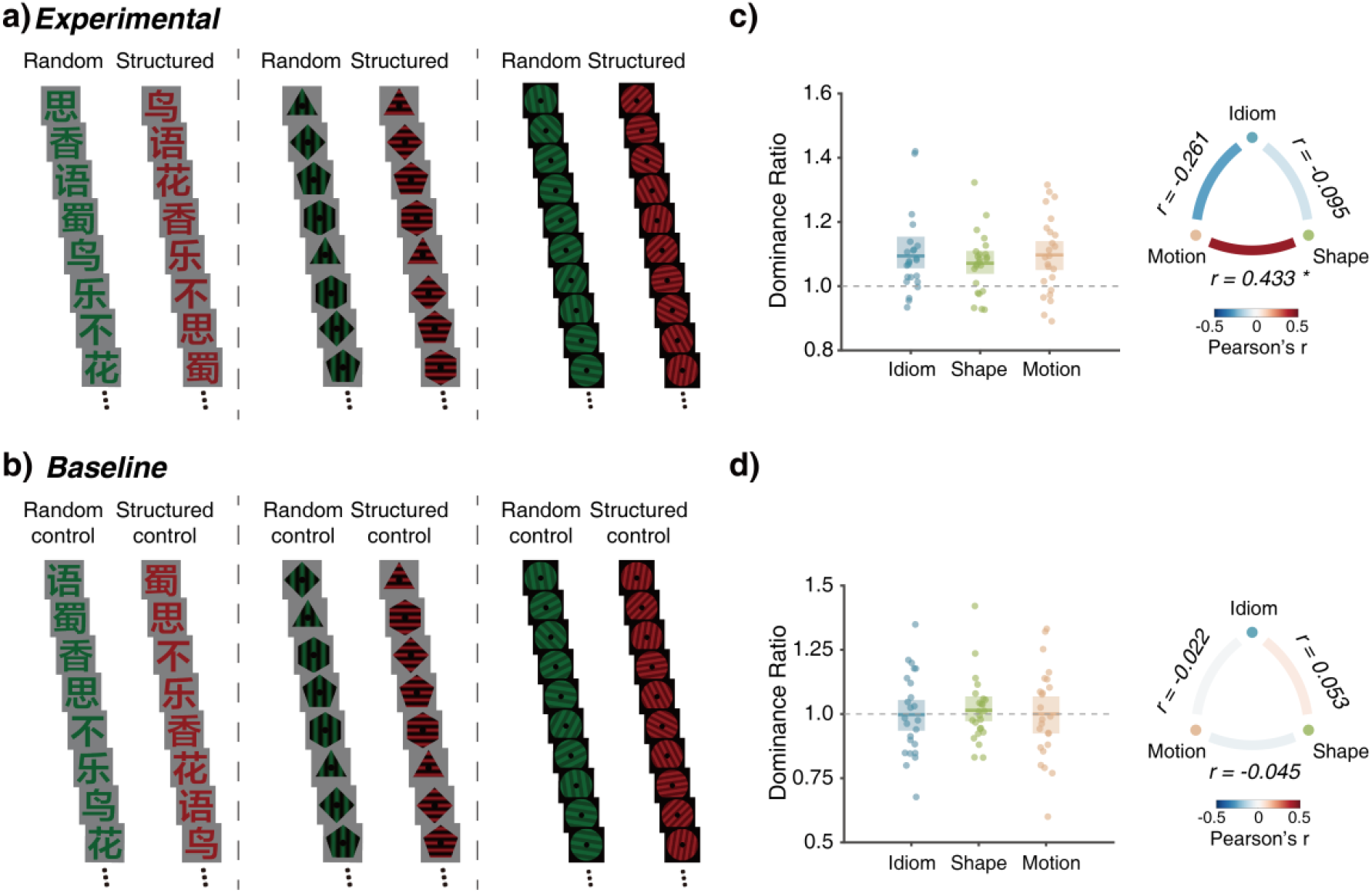
Example stimuli and results of Experiment 4. a) In the experimental conditions, the stimuli were the same as that in Experiment 1, except that for shape and motion, different spatial frequencies were assigned to the structured and random streams. b) In the baseline conditions, the structured-control streams were obtained by temporally reversing the sequence order of the corresponding structured streams (for Chinese idioms), or by randomizing the stimulus order of each rhythmic cycle, i.e., every four items (for motion and shape), while retaining other stimulus properties. The random-control streams were generated in the same way as that for the random streams in the experimental condition. Note that for the shape and motion stimuli, the structured-control and random-control streams had the same spatial frequencies as their corresponding counterparts in the experimental conditions. c) & d) Dominance ratios of the three stimulus types and the pairwise Pearson correlation results, for the experimental conditions and the baseline conditions. The colored lines and bars represent the averaged dominance ratios with bootstrapped 95% CIs, and each colored dot represents one participant. * p < 0.05; **: p < 0.01.

Results from the experimental condition replicated that from Experiment 1 as all three types of structured streams held a perceptual advantage over their random counterparts (Fig. 3c; Idiom: *t* (23) = 3.80, *p* < 0.001, Cohen’ *d* = 0.77; Shape: *t* (23) = 3.75, *p* < 0.001, Cohen’ *d* = 0.76; Motion: *t* (23) = 3.93, *p* < 0.001, Cohen’ *d* = 0.80). Moreover, Pearson correlation analysis of dominance ratios yielded a significant correlation only between the shape and motion conditions (*r* = 0.43, *p* = 0.03), but not between idiom and shape (*r* = -0.26, *p* = 0.21) or between idiom and motion conditions (*r* = -0.09, *p* = 0.68). For the baseline conditions, by contrast, the benefit in BR was eliminated for the structured-control streams, in all three stimulus types (Fig. 3d; *ps* > 0.5; Idiom: Cohen’ *d* = 0.01; Shape: Cohen’ *d* = 0.15; Motion: Cohen’ *d* = 0.01). In addition, there was no reliable correlation between any of the stimulus pairs (*ps* >= 0.8).

More importantly, for each stimulus type, we evaluated whether the structured streams gained perceptual advantages from the dominance (conscious) or the suppression (nonconscious) period of BR. During the dominance phase (Fig. 4a), the mean normalized duration was significantly prolonged for the structured streams relative to the corresponding baseline for the idiom condition (*t* (23) = 2.30, *p* = 0.03, Cohen’ *d* = 0.47), but not for the shape (*t* (23) = 0.26, *p* = 0.79, Cohen’ *d* = 0.05) or the motion condition (*t* (23) = 1.22, *p* = 0.23, Cohen’ *d* = 0.25), indicating a privilege of regular structure at the conscious level restricted to semantic-level regularities. During the suppression phase of the structured information (Fig. 4a), on the other hand, we found significantly reduced suppression durations for the structured streams as compared with the baseline for both the shape (*t* (23) = 2.01, *p* = 0.05, Cohen’ *d* = 0.41) and motion conditions (*t* (23) = 2.15, *p* = 0.04, Cohen’ *d* = 0.44), but not for the idiom condition (*t* (23) = 0.15, *p* = 0.87, Cohen’ *d* = 0.03), suggesting an advantage at the nonconscious level induced by perceptual-level regularities.

**Fig. 4.**
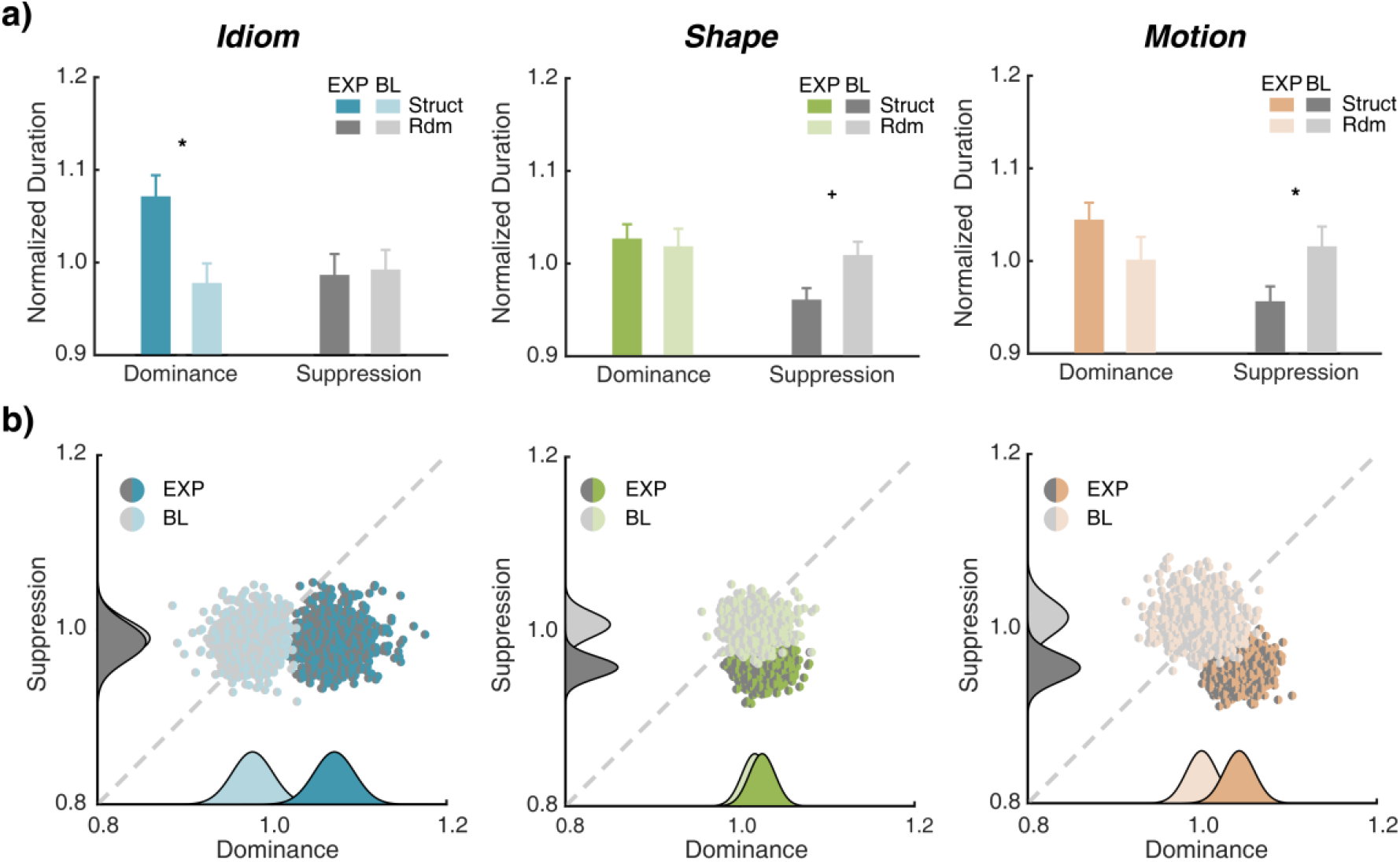
Conscious and nonconscious benefits for semantically and perceptually defined temporal structures during the BR. a) Normalized mean percept durations of structured and structured-control streams during their dominance phase (color bars) and suppression phase (grey bars, when random and random-control streams were in dominance). Error bars: SE. * *p* < 0.05, + *p* = 0.05. b) Bootstrap distributions of mean normalized dominance durations for each stimulus. The bootstrapped data for the dominance and suppression phases of the structured stream was projected onto the x and y axes, respectively, with the projected data fitted using a Gaussian probabilistic density function (PDF). In each dimension (i.e., on the x or the y axis), the evaluated values of the 2 PDFs were pooled together and then remapped into an appropriate range by applying a linear transformation for visualization purposes. These fitted curves showed a clear dissociation of the idiom and the other stimuli along the conscious and nonconscious dimensions.

For better comparison within each stimulus condition, we visualized the bootstrap distributions of normalized durations in the experimental and the baseline conditions based on one thousand resampled data sets (Fig. 4b). The horizontal and vertical axes represent the conscious (dominance) and the nonconscious (suppression) phases of the structured/structured-control stream. It is clearly shown that, for the idiom stimuli, the resampled data sets separate in the dominance rather than the suppression dimension, which means the observed privilege of structured idiom streams arises mainly from lengthened perception when they dominate conscious awareness. By contrast, for the shape and motion streams, the experimental and baseline conditions diverge primarily at the suppression phase, highlighting the contribution from nonconscious processing of the structured information.

### Privilege of the perceptual-level temporal structures in conscious access

The outcomes of BR reflect a combined effect of conscious and nonconscious visual processing. Some may argue that the lack of nonconscious privilege of the structured idiom stream in Experiment 4 may result from the disturbance of the conscious processes. To solve this problem, in Experiment 5, we directly probed the role of regular temporal structures in nonconscious visual processing using the breaking continuous flash suppression (b-CFS) paradigm (Fang and He, 2005; Tsuchiya and Koch, 2005). By presenting a salient dynamic noise to one eye and the target stimulus to the other, we created a situation where the target was continuously suppressed until it broke into awareness (Fig. 5a). The time for the target to reach awareness can serve as an index to assess the potential effects associated with nonconscious visual perception (Jiang et al., 2007).

**Fig. 5.**
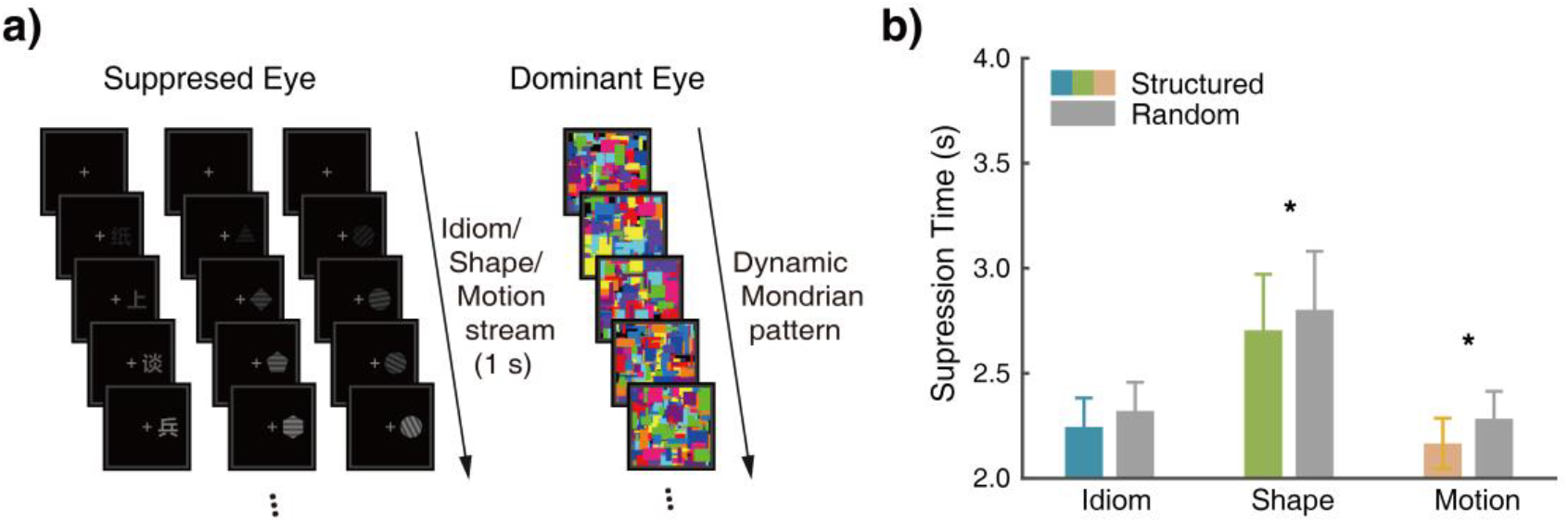
Schematics of the stimulus sequences used in the b-CFS Experiment and the suppression time results. a) A dynamic Mondrian pattern was presented to the observer’s dominant eye to suppress a structured or a random stream presented to the non-dominant eye from the beginning of each trial. The contrast of the stimulus stream was ramped up from 0 to an individually adjusted value within 1-2 s and stayed constant thereafter. Participants were instructed to indicate on which side of the fixation the target appeared, and their reaction times (i.e., suppression times) for each stimulus condition were shown in b). Error bars: SE. * *p* < 0.05.

Paired sample t-tests revealed that for the shape and motion conditions, the perceptual-level structured streams emerged from suppression significantly faster than the random ones (Fig. 5b, Shape: *t* (17) = 2.29, *p* = 0.03, Cohen’s *d* = 0.54; Motion: *t* (17) = 2.62, *p* = .01, Cohen’s *d* = 0.61); whereas for the idiom condition, the difference of suppression time between the structured and random streams was not significant (*t* (17) = 1.37, *p* = 0.19, Cohen’s *d* = 0.32). These findings provide substantial evidence that perceptual-level regularities in temporal structures of visual information could be extracted without awareness while semantic-level structures could not, which is in good accordance with the dissociation of conscious and nonconscious privileges of temporal structures observed in the BR experiment.

## Discussion

### A multi-level privilege of regular temporal structure in visual competition

By adopting dynamic visual streams that are temporally organized based on certain regularities, our study provides novel evidence that during the interocular competition, visual awareness is biased towards information with regular temporal structures. Specifically, we found structured streams formed by several types of visual stimuli, including Chinese idiom, shape, and motion, enjoyed longer dominance durations relative to their random counterparts in BR. Such an advantage is not attributable to low-level factors, as the rivalry streams always consisted of the same elements though being arranged in different sequences. Nor can it be accounted for solely by the repetitions of physical stimuli, given that the effect persisted even for structured streams constructed by non-repeated Chinese idioms. Alternatively, the current findings highlight a factor broadly related to information structure in the temporal dimension that can robustly modulate visual competition across different stimulus levels.

These findings, however, do not warrant that the prioritization of regularity in dynamic visual competition is a universal rule that applies to all types of visual stimuli. Indeed, the perceptual advantage induced by perceptual-level regularities did not extend to rhythmic structures defined by contrast change. The difference between contrast and the other stimulus conditions probably lies in that contrast is among the most basic visual features that can be well resolved by neuronal responses within the primary visual cortex (Goodyear and Menon, 1998), which makes contrast perception more vulnerable to changes in stimulus strength. As stated in Levelt’s proposition about BR, increasing stimulus strength for one eye or its relative strength to the other eye will enhance the perceptual predominance of the corresponding stimulus (Levelt, 1965). Accordingly, the successive change in relative contrast strength between the two eyes’ stimuli may cause stochastic fluctuations in their relative predominance during BR, thereby abolishing the perceptual advantage for the structured contrast streams. Although this assumption remains to be verified, our findings have set constraints on the level at which temporal structures modulate visual competition, suggesting a mechanism that probably operates beyond the early (monocular) stage of visual processing.

Despite the lack of effect in the contrast condition, what gives rise to the observed advantage of the structured shape, motion, and idiom streams? Does a common mechanism underlie the advantage shared across these stimulus types? The inter-stimulus correlation analysis provided an initial hint on these issues: significant correlations of the dominance ratio were found only between the motion and shape, but not between the idiom and the other two stimulus conditions, indicating that perceptual- and semantic-level regularities might not modulate perceptual awareness via a unified mechanism. A more powerful way to test this proposition is to identify the boundary conditions for the observed effect with different stimuli. To this end, we manipulated the spatiotemporal integration window of the rivalry streams (Experiments 2 & 3). When removing spatial or temporal uniformity from the rivalry streams, the perceptual advantage of structured streams was still significant in the idiom condition but disappeared for the motion and shape conditions, suggesting a dissociation between the effects caused by semantic- and perceptual-level regularities. Intriguingly, such dissociation was also manifested in the results that structured streams defined by two types of regularities modulated visual competition at different conscious stages (Experiments 4 & 5). On the one hand, temporal structures built on rhythmic changes of physical features held an advantage largely stemmed from nonconscious visual integration, as indicated by reduced durations in the suppression phase of BR as well as the more rapid breakthrough from CFS than the random streams. On the other hand, the advantage of temporal structure emerging from semantic-level regularities originated mainly from conscious information integration, as evidenced by prolonged perceptual durations during the dominance phase of BR and the lack of advantage in the CFS experiment. Collectively, these results suggest that the privileges of perceptual- and semantic-level regularities are unlikely to be driven by a simple common mechanism, albeit there is a general tendency for the human brain to prioritize multi-level structured information during visual competition.

### Information integration and consciousness

The relationship between integration and awareness is central to contemporary accounts of consciousness (Dehaene and Naccache, 2001; Lamme, 2018; Mudrik et al., 2014; Tononi et al., 2016), as it lays a theoretical foundation for characterizing the physical substrates of consciousness (Tononi et al., 2016), and provides critical insights into the neural and computational models of visual awareness (Mashour et al., 2020; Oizumi et al., 2014). According to the integrated information theory (IIT), the content of consciousness can be qualitatively identified with and quantitatively measured by the process of information integration. In particular, a conscious experience is identical to a conceptual structure encompassing a maximum of integrated information, with the extent of integration to be specified by the variable Φ^max^ (Tononi et al., 2016). In this view, the observed perceptual advantage of structured information streams can be accounted for by enhanced temporal information integration. The integration of visual elements over time may benefit from their intrinsic connections defined by certain types of regularities, allowing the incorporation of discrete events into a conceptual structure with a higher value of Φ ^max^ such that facilitates the emergence of subjective conscious experience (Oizumi et al., 2014; Tononi et al., 2016).

The IIT would predict an overall advantage of structured streams during the visual competition. However, it is insufficient to explain the different boundary conditions we found between semantic- and perceptual-level stimuli regarding the temporal integration process, especially their links with conscious and nonconscious processing, respectively. It has long been assumed that consciousness is necessary for information integration to take place (Dehaene and Changeux, 2011; Mashour et al., 2020). Nonetheless, recent studies reveal that some integration processes, ranging from the integration of static natural scenes (Mudrik et al., 2011a), dynamic motion signals (Faivre and Koch, 2014) to that of symbolic information based on simple semantic or arithmetic relations (Sklar et al., 2012; Yeh et al., 2012), and even multisensory information (Alsius and Munhall, 2013) can occur without awareness. These findings stress the need to specify the function of consciousness at different levels of integration. In the current study, by directly comparing various types of stimuli with the same (BR and bCFS) paradigms, we identified the limits of nonconscious visual processing from the aspect of temporal integration. Firstly, we found regular shape and motion streams but not idiom streams could enjoy a perceptual privilege even below consciousness. These observations coincide with previous findings of nonconscious adaption to invisible apparent motion and biological motion sequences (Faivre and Koch, 2014), suggesting that consciousness may not be required for the temporal integration of certain types of rhythmically changed visual patterns. On the other hand, empirical results concerning the integrative processing of invisible semantic information seem to be more complicated. Using the semantic priming paradigm, some studies have reported evidence for nonconscious temporal integration of meaningful contents based on relatively simple semantic relations (Yeh et al., 2012), while others failed to find such effect for sequentially presented characters within an idiomatic context (van Gaal et al., 2014; Zhou et al., 2016). Using the BR paradigm, one study showed that audiovisual interaction could enhance the temporal integration and boost the visual awareness of musical scores, but only by lengthening the dominance (i.e., conscious) phase (Lee et al., 2015). In conjunction with these studies, our finding that the perceptual advantage of concatenated idiom streams transpired primarily above the conscious level suggests that consciousness is necessary at least for the temporal integration of symbolic information based on complex rules, which enriches our understanding of the notion that “consciousness is needed when the integration exceeds a certain level of complexity” (Mudrik et al., 2014).

The dissociation in conscious demands among different stimulus streams is also compatible with our finding that the privilege of perceptual-but not semantic-level information is vulnerable to the lack of spatiotemporal uniformity in the information integration window. Compared with conscious integration, nonconscious integration is limited by relatively smaller temporal integration windows and simpler associations (Armstrong and Dienes, 2013; Faivre and Koch, 2014). This difference may explain why increasing the variance in temporal durations or spatial locations of individual items only have detrimental effects for structured shape and motion streams (Experiments 2 & 3), as their benefits originate mainly from nonconscious processing.

### Possible mechanisms underlying the temporal structure privilege

One might argue that the temporal structure advantage observed in the current study can be adequately explained by the attentional processes. Research has shown that top-down or endogenous attention can bias perceptual dominance to some degree during binocular rivalry (Chong et al., 2005; Meng and Tong, 2004). Nonetheless, there are several reasons that these attentional modulations may not cause the effects obtained from our study. Firstly, without noticing the regular structure of the monocular stream, observers were not able to track such temporal structure through voluntary control. Indeed, according to the post-debriefing assessment in Experiment 1, none of the participants were aware of the existence of regular structure in the shape, motion, and contrast conditions. Moreover, previous research suggests that voluntary attention can mediate perceptual dominance by decreasing the mean dominance durations of the non-attended stimulus (Hancock and Andrews, 2007). Inconsistent with this explanation, we observed prolonged dominance durations for the structured idiom streams rather than decreased durations for the random ones. On the other hand, the effect of exogenous attention in BR has been evidenced by enhanced initial dominance or prolonged dominance durations due to a difficulty to disengage attention from a visual object (Mitchell et al., 2004) or an object incongruent with the semantic context (Mudrik et al., 2011b; Wolf and Hochstein, 2011). However, such object-based or violation-related attentional bias could neither account for the perceptual advantage that we observed. Above all, an additional data analysis revealed no evidence for the initial dominance bias predicted by object-based attention (see Supplementary Information). Besides, previous studies showed a greater predominance of contextually incongruent objects during BR, while here, the ‘congruent’ stimuli (i.e., structured streams) did dominate participants’ subjective experience for a longer time. Taken together, the current findings are not likely to be directly driven by the attentional effect, although they do not negate the general contribution of attention to BR. Future research could try to examine the roles of temporal structure on visual attention and awareness, respectively, given the intricate relationship between these two phenomena (Cohen et al., 2012).

More importantly, our findings provide fresh insight into the controversial issue regarding how meaningful stimulus content modulates perceptual predominance in visual competitions. Using static images, previous studies have demonstrated a variety of modulatory effects in BR that can be ascribed to a general factor, namely, the ‘meaningful content’ of the stimulus (Walker, 1975). Nonetheless, the influence of such effects could be in either direction (promoting or repressing), as demonstrated by research on both the figure (Yu and Blake, 1992) and word competitions (Wolf and Hochstein, 2011). The discrepancy in these results raises a fundamental question: whether the effect of ‘meaningful content’ actually arises from several distinct mechanisms. As aforementioned, an attentional influence may explain the observed disadvantage of a meaningful object when competing with a meaningless (yet structured) one (Mudrik et al., 2011b). What remains open, however, is the cause for the opposite effect, namely, the perceptual advantage observed for relatively more meaningful content. The structured streams used in our study are arguably more meaningful than the random ones and enjoy a privilege in BR, regardless of the types of regularities. These findings resemble the advantage of recognizable figures vs. scrambled images and statically rendered idioms vs. non-idioms (Wolf and Hochstein, 2011; Yu and Blake, 1992). Notably, the meaningful stimuli in these studies (including ours) all have an intrinsic structure that can be captured through the integration of visual elements, while the meaningless stimuli do not. Therefore, it is tempting to speculate that the superiority of meaningfulness in visual competition is mediated, at least in these cases, by mechanisms facilitating information integration. Regularities in information structure, either in the dimension of space or time, may provide a predictive context for the brain to more easily organize discrete pieces of information into a coherent conscious experience, and further bias the content of awareness in a situation with uncertainty such as the BR (Hohwy et al., 2008). This notion offers a promising account for the perceptual advantage of a diversity of ‘meaningful’ (or rather structured) stimuli in the visual competition, opening up a new avenue worthy of further exploration.

### Neural basis for the multi-level temporal structure privilege

BR is driven by reciprocal inhibitions of the binocular signals in the early visual areas (Leopold and Logothetis, 1996; Polonsky et al., 2000), as well as frontoparietal signals that may trigger a reorganization of activities in early sensory areas through feedback (Knapen et al., 2011; Lumer et al., 1998), although the role of frontal activity in directly mediating spontaneous perceptual alternations is still under debate (Brascamp et al., 2018; Frassle et al., 2014). Among these areas, in which locus the privilege of structured information arises remains an open issue. However, it is plausible to assume that the possible locus may receive modulatory signals from stimulus-specific brain regions actively involved in visual temporal integration and temporal structure extraction. We conjecture these regions may reside in higher levels of the visual hierarchy (beyond the monocular stage), considering the lack of perceptual advantage for the regular contrast sequences, and include the language processing network specifically for the idiom condition.

From the perspective of temporal coding, the privilege of structured streams is established gradually during the rivalry rather than upon immediate exposure to the rivalrous stimuli. How does such advantage arise in the brain? Likely, the abovementioned interaction between brain regions in charge of the temporal structure processing and visual competition is implemented dynamically through neural entrainment, a phenomenon that the intrinsic neural oscillations synchronized to the rhythms of external events (Haegens and Zion Golumbic, 2018; Thut et al., 2011). Neural entrainment is critically involved in the perception of simple (e.g., tone sequences) and complex (e.g., music and language) auditory streams, as well as in the processing of rhythmic visual information (Brookshire et al., 2017; Haegens and Zion Golumbic, 2018). Entraining neural oscillations at different brain areas to the same external rhythm (such as the structure-derived delta rhythm in the current study) may enhance neural communications within the coactivated regions (Fiebelkorn and Kastner, 2020), so that strengthen the neural representation of the structured information to bias visual awareness. Future studies are needed to elucidate the exact mechanisms and the neural circuits underlying the emergence of dynamic conscious contents based on multi-level temporal structures.

## Conclusion

To recapitulate, we show that during the dynamic visual competition, the content of visual awareness is biased towards information with regular temporal structures. While such advantages can be generalized to semantic- and perceptual-level regularities, the underlying benefits respectively stem from conscious and nonconscious temporal integration, suggesting that they are mediated by partially overlapped but distinct mechanisms. These findings corroborate and extend the IIT by highlighting the crucial role of structure-guided information integration in the continuous emergence of conscious content. They also shed new light on the diversified functions of subjective awareness in temporal integration across the hierarchy of visual information processing.

## Methods

### Participants

A total of one-hundred and fifty native Chinese speakers (mean age ± SD = 22.4 ± 2.7 years, 78 females) took part in this study. Seventy-two participated in Experiment 1 (18 in each stimulus condition), 18 in Experiments 2, 3, and 5, respectively, and 24 in Experiment 4. All participants had normal or corrected-to-normal vision with no reported history of strabismus, and exhibited a rivalry pattern without extreme eye/color dominance (less than 90%) or a large proportion of mixture state (less than 30% of the tracking period) during a pre-screening test. All participants were naïve to the purpose of the experiment. They gave informed consent to participate in procedures approved by the institutional review board of the Institute of Psychology, Chinese Academy of Sciences.

### Apparatus and stimuli

Visual stimuli were generated using MATLAB (The MathWorks, Natick, MA) with the Psychtoolbox extension (Brainard, 1997; Pelli, 1997), and displayed on a 21-inch CRT monitor with a resolution of 1280 × 1024 at 60 Hz. Observers viewed the stimuli through a mirror stereoscope at a distance of 60 cm, with the head stabilized by a chin rest.

In the BR experiments, the rivalry streams were tinted in red and green with matched luminance values (4.87 cd/m^2^), and displayed on a black (or gray for the idiom condition) background to ensure that the participants achieved a relatively stable rivalry performance. Each pair of the rivalry streams consisted of the same 120 items but with different orders to form structured and random sequences. Four types of stimulus streams, including idiom, shape, motion, and contrast (for Experiment 1 only), were employed.

For the idiom condition, we randomly selected 30 different four-character Chinese idioms from a pool of 60 idioms (familiarity > 6.4 on a 7-point scale) and concatenated them to form a structured stream for each trial. To obtain a random counterpart, we shuffled the structured stream at the character level so as to eliminate the semantic structure.

For the shape condition, the structured streams were composed of Gabor patches embedded in polygonal contours. The edge number of the shape increased monotonically every four items (i.e., triangle ⟶ diamond ⟶ pentagon ⟶ hexagon), yielding a rhythmic pattern throughout the trial. The random streams were generated by randomizing the sequence order within each rhythmic cycle (i.e., every four items) of the structured streams with the constraint that not any two successive items were the same. The orientations of the Gabor patches for each rivalry pair were set to horizontal and vertical respectively. Each polygonal shape subtended 2.42° of visual angle in terms of circumradius with a spatial frequency of 2.4 cycles per degree (cpd) for Experiments 1-3, and 2.4 or 3.3 cpd for Experiment 4. Both the structured and random streams were superposed on a gray square to facilitate the perception of the edges.

For the motion condition, the structured streams consisted of a grating (radius: 2.42°; spatial frequency: 2.4 cpd or 3.3 cpd (for Experiment 4 only); initial orientation set at -45° or 45° from vertical) rotating clockwise in step of 36°. Thus, every ten steps of movements formed one cycle of the rhythmic structure (i.e., a full circle of 360°). The random streams were generated in the same way as in the shape condition.

For the contrast condition, the structured stream consisted of a grating whose Michelson contrast increased from 0.4 to 0.7 (step = 0.1) in each rhythmic cycle. The random streams were generated in the same way as in the shape condition. In each trial, the paired gratings were tilted -45° and 45° from vertical respectively, with the other parameters identical to that for the motion condition.

In the b-CFS experiment, a dynamic Mondrian pattern composed of partly overlapping rectangles of varying sizes and colors was presented to the observer’s one eye, which suppressed the awareness of the structured/random stream (2.12° × 2.12°) presented to the other eye. The suppressed streams were rendered in gray against a black background but otherwise the same as that used in Experiment 1.

### Procedures

Prior to each experiment, participants were instructed to adjust the mirror stereoscope in order to achieve successful binocular fusion. In Experiment 1, participants were randomly assigned to one of the four stimulus conditions (Fig. 1a). They started with a practice session to get familiar with the stimuli and procedures, followed by one test session divided into two blocks. Each block consisted of 8 trials, with the color assignment, the eye, and the stimulus orientation (for the shape and contrast conditions only) counterbalanced across trials. Each trial began with a central fixation dot presented to both eyes. Participants were instructed to maintain fixation on this point throughout the whole trial. After two seconds, the structured and random streams were presented dichoptically in synchronization, 250 ms for each item with no intervals, during the 30 s rivalry period. Participants were required to press and hold one of two keys to indicate which color (red or green) of the tinted streams was dominating perception, and to release keys when perceiving a fused plaid or a piecemeal rivalry. To avoid potential fatigue effects due to the interocular competition, each trial was followed by a compulsory break (inter-trial interval, ITI) of 8 s.

In Experiments 2 and 3, the procedure was the same as that for Experiment 1, but with the following exceptions. In Experiment 2, the duration of each individual item was no longer constant (250 ms) but changed irregularly with an up to 50% variance (ranging from 125 to 375 ms in step of 16.7 ms), thereby disrupting the regularities of stimulation in time. In Experiment 3, the location of each stimulus was no longer fixed at the retinal center but altered randomly within a range of 0° to 1.51° from the center, thus destroying the regularities of stimulation in space. Moreover, in both Experiments 2 and 3, each participant completed two rivalry blocks (a total of 16 trials) for each of the three stimulus conditions (i.e., idiom, shape, and motion). The order of stimulus conditions was counterbalanced across participants.

In Experiment 4 (Fig. 3), we replicated Experiment 1 except for the contrast condition. Critically, for each stimulus type, we added a baseline condition where the two rivalry streams were both deprived of regularity-defined temporal structure, but correspondingly matched with the structured and random streams in the experimental condition to control for low-level properties. For idiom, the structured-control streams were obtained by temporally reversing the sequence order of the structured idiom stream, which removed the ruled-based semantic relationships from each idiom while retaining the cooccurrence-based association among its four characters. For shape and motion, the structured-control streams could not be created by simply reversing the structured streams, because the temporal structure would remain regular after temporal reversal. Therefore, we randomized the stimulus order within each rhythmic cycle (every four items) of the structured shape or motion stream to remove the perceptual-level regularities. Meanwhile, we set different spatial frequencies for the Gabor patches that constituted the rivalry streams, and matched the spatial frequency of the stimuli between the baseline and experimental conditions (3.30 cpd for structured and structured-control streams and 2.40 cpd for random and random-control streams or vice versa). The spatial frequency of the rivalry streams and the order of stimulus conditions were counterbalanced with a Latin square design across participants. Each participant completed 3 BR blocks, each for one stimulus condition. Each block consisted of 16 trials, with the baseline and experimental trials presented in a pseudo-randomized sequence (‘ABBABAAB’ or vice versa for every 8 trials).

In Experiment 5, participants started with a practice session to get familiar with the stimuli and procedures, followed by one test session consisting of 90 trials. Among these trials, 40 were structured trials, another 40 were random trials, and the remaining 10 were catch trials in which no stimuli appeared in the suppressed eye. In each trial, participants were asked to maintain fixation on a central cross that was continuously presented to both eyes. To ensure that the stimulus stream could be adequately processed below awareness with sufficient time, a dynamic noise pattern was presented to the dominant eye at full contrast, and a structured or random stream was presented to the other eye. The stimulus center was located 1.86° to the left or the right of the fixation. Moreover, the contrast of the stimulus stream was ramped up gradually from 0 to a value between 0.2 and 1 within a limited duration (1 or 2 s), with the values determined based on individual performance during the practice session. After that, the stimulus contrast remained constant until a response was made or 10 s elapsed. Participants were instructed to press the corresponding button to indicate on which side the target appeared.

### Data Analysis

In Experiments 1–4, to obtain reliable estimates of the temporal dynamics of binocular rivalry, we first identified and removed the unstable percepts based on the following criteria. Button press durations less than 200 ms were not counted as periods of exclusive dominance. Besides, during transitions from pressing one button to the other, participants sometimes accidentally pressed both buttons for short periods (< 200 ms). Such responses were also excluded from further analysis. Based on the remaining responses, we computed the trial-based accumulated dominance duration (i.e., the overall dominance duration of a given percept within each 30 s trial averaged across all trials) and the percept-based mean dominance duration (i.e., the dominance duration of a given percept in all trials averaged across all percepts) for the structured and the random streams respectively, for each stimulus type and each participant. To facilitate comparison among different stimulus types, we calculated the dominance ratios of the structured to the random stream based on the accumulated and mean dominance durations. In Experiment 1, we observed similar effects with the two dominance ratio indices for all stimulus types, so in Experiments 2-4, we only reported results based on the mean dominance ratios. Additional analysis based on the accumulated dominance ratios yielded very similar patterns, which were reported in the supplementary file.

To obtain a more robust estimate of the confidence intervals than the standard methods, we used a standard bootstrap procedure (*n* = 1000) to compute the bias-corrected and accelerated 95% confidence intervals (CI) for the average of these dominance ratios (Davison and Hinkley, 1997; DiCiccio and Efron, 1996). The bootstrap procedure was also applied to the normalized dominance durations of the structured and random streams in Experiments 1 and 4 to illustrate the reliability of the observed difference among stimulus conditions. In Experiment 1, the normalization was carried out through dividing the mean dominance durations of the structured and the random streams respectively by the average of dominance durations of all percepts (across both streams) for each participant. In Experiment 4, the dominance durations of structured and random streams were divided respectively by the individual mean of overall dominance durations across the experimental and baseline conditions for each stimulus.

In Experiment 5, we measured the suppression time as the reaction times (RTs) needed for the participants to correctly indicate at which side of the fixation they saw the target stimulus. For each individual, only trials with RTs within three times of the standard deviation of the individual mean based on all trials were included in further analysis.

## Acknowledgements

We are grateful to Randolph Blake for his inspiring comments and discussion about some preliminary findings from the current study. This research was supported by grants from the National Natural Science Foundation of China (Nos. 31525011, 31771211, and 31830037), the Strategic Priority Research Program (No. XDB32010300), the Key Research Program of Frontier Sciences (No. QYZDB-SSW-SMC030), the Youth Innovation Promotion Association of the Chinese Academy of Sciences, the National Key Research and Development Project (2020AAA0105600), Beijing Municipal Science & Technology Commission, Shenzhen-Hong Kong Institute of Brain Science, and the Fundamental Research Funds for the Central Universities.

