## Supplemental Information for "Visual temporal integration by multi-level regularities fosters the emergence of dynamic conscious experience"

### Supplementary Information

#### Analysis of accumulated dominance ratios

In Experiments 2 and 3, the dominance ratios based on accumulated dominance durations of structured and random streams (Table S1) were significantly greater than 1 for the idiom condition (Experiment 2: $t$ (17) = 3.11, $p$ < 0.01, Cohen' $d$ = 0.73; Experiment 3: $t$ (17) = 2.51, $p$ = 0.02, Cohen' $d$ = 0.59). Whereas for the shape condition, there was no evident advantage of structured streams over the random ones (Experiment 2: $t$ (17) = 0.23, $p$ = 0.81, Cohen' $d$ = 0.01; Experiment 3: $t$ (17) = 0.00, $p$ = 0.99, Cohen' $d$ = 0.00), and similar pattern was observed in the motion condition (Experiment 2: $t$ (17) = 1.09, p = 0.28, Cohen' d = 0.25; Experiment 3: $t$ (17) = 0.23, $p$ = 0.81, Cohen' $d$ = 0.05).

In Experiment 4, additional analyses on accumulated dominance ratios (Table S1) revealed that all three types of structured streams held a perceptual advantage over the random counterparts in the experimental conditions (Idiom: $t$ (23) = 3.53, $p$ < 0.01, Cohen' $d$ = 0.72; Shape: $t$ (23) = 3.03, $p$ < 0.01, Cohen' $d$ = 0.61; Motion: $t$ (23) = 3.29, $p$ < 0.01, Cohen' $d$ = 0.67), but not in the baseline conditions (Idiom: $t$ (23) = 0.71, $p$ = 0.48, Cohen' $d$ = 0.14; Shape: $t$ (23) = 0.09, $p$ = 0.92, Cohen' $d$ = 0.01; Motion: $t$(23) = 0.35, $p$ = 0.72, Cohen' $d$ = 0.07).

**Table S1 The accumulated dominance ratio results from Experiments 2 – 4**

| | Exp. 2 | Exp. 3 | Exp. 4 - EXP | Exp. 4 - BL |
|---|---|---|---|---|
| | *M (SD)* | *M (SD)* | *M (SD)* | *M (SD)* |
| Idiom | 1.07 (0.10) | 1.05 (0.09) | 1.09 (0.13) | 1.02 (0.17) |
| Shape | 1.00 (0.07) | 1.00 (0.13) | 1.05 (0.09) | 1.00 (0.12) |
| Motion | 1.02 (0.09) | 1.00(0.07) | 1.11 (0.16) | 1.01 (0.21) |

*Note. M* and *SD* represent mean and standard deviation, respectively.

**Analysis of initial dominance indices**

An additional initial dominance analysis was performed based on the data obtained from Experiment 1 to confirm that the advantage of structured information was not due to the effect of object-based attention, which could be established promptly at the initial stage of each trial. To this end, we calculated the proportion of trials in which the structured stream dominate perception at the beginning of the trial (i.e., the 'onset dominance'). Besides, we also calculated the proportions of trials in which participants saw the structured stream after the first cycle of temporal structure (i.e., the 'post-exposure dominance'). Analysis on both measures provide no evidence for an initial attentional bias. Firstly, the 'onset dominance' proportions for idiom, shape, motion, and contrast were 51%, 45%, 47%, and 48%, respectively, none of them significantly differed from 50% (Idiom: $t$ (17) = 0.59, $p$ = 0.56; Shape: $t$ (17) = 1.58, $p$ = 0.13; Motion: $t$ (17) = 1.63, $p$ = 0.12; Contrast: $t$ (17) = 0.65, $p$ = 0.52). Moreover, similar results were found on the 'post-exposure dominance' proportions, which were 52%, 46%, 46%, and 49%, respectively, and not significantly different from 50% either (Idiom: $t$ (17) = 0.84, $p$ = 0.40; Shape: $t$ (17) = 0.91, $p$ = 0.37; Motion: $t$ (17) = 1.36, $p$ = 0.19; Contrast: $t$ (17) = 0.32, $p$ = 0.74).
